# A Quantitative Study of Inappropriate Image Duplication in the Journal *Toxicology Reports*

**DOI:** 10.1101/2023.09.03.556099

**Authors:** Sholto David

## Abstract

Inappropriate image duplication is a type of scientific error that can be detected by examining published literature. Few estimates of the frequency of this problem have been published. This study aimed to quantify the rate of image duplication in the journal *Toxicology Reports*. In total 1540 unique articles (identified by DOI) were checked for the presence of research related images (microscopy, photography, western blot scans, etc). Each research paper containing at least one such image was scrutinized for the presence of inappropriate duplications, first by manual review only, and subsequently with the assistance of an AI tool (ImageTwin.ai). Overall, *Toxicology Reports* published 715 papers containing relevant images, and 115 of these papers contained inappropriate duplications (16%). Screening papers with the use of ImageTwin.ai increased the number of inappropriate duplications detected, with 41 of the 115 being missed during the manual screen and subsequently detected with the aid of the software. In summary, the rate of inappropriate image duplication in this journal has been quantified at 16%, most of these errors could have been detected at peer review by careful reading of the paper and related literature. The use of ImageTwin.ai was able to increase the number of detected problematic duplications.

## Introduction

Scientific papers often include images to communicate the results of experiments or illustrate case reports. Mislabelled images mislead the reader of a paper (intentionally or not), and when discovered, undermine the credibility of research. One way that incorrectly labelled images can be discovered is by identifying duplicated images – where one image is used to represent more than one experimental condition or subject, or parts of an image are cloned to produce composite images. The frequency of image duplication in biomedical literature has previously been estimated at ∼5%^1,2^. Data duplication between papers, including charts as well as images, has been quantified as high as ∼25%^3^. Greater awareness of the problem of image duplication has led to increased documentation of such errors, often on the post-publication peer review website PubPeer^4^. Recently, AI tools have become increasingly capable in detecting image similarities, with several tools being developed specifically for detecting image manipulation or duplication in the scientific literature. A recent study using the software ImageTwin.ai reported that 9 out of 69 rhinology related papers screened using the software contained inappropriate image duplication^5^. The purpose of this study was to quantify the rate of inappropriate image duplication in the open access journal *Toxicology Reports* and compare the results of manually screening papers to a review assisted by the ImageTwin.ai tool.

## Methods

### Paper Selection

All research articles or case reports published in the journal *Toxicology Reports* (since the first article in 2014 to the most recent article at the time of writing in July 2023) were screened for the presence of any images used to illustrate or communicate the results of experiments, typically histology slides, microscopy images of cells, photographs of animals, western blots, and DNA gels. Articles containing only charts, spectra, maps, graphical abstracts, diagrams, or computer-generated images were excluded. Review articles, retraction notes, letters, and editorials were excluded as they are not expected to contain original images.

### Manual Screen of Papers for Inappropriate Duplication

The PDF version of each included article was scrutinized by one reviewer (the author) for evidence of duplication within each paper. Supplementary data files were also occasionally analysed for inappropriate duplications, but not systematically (some sets of papers were bulk downloaded as PDFs without visiting the journal page for each article). In some cases, images were also compared between other papers by the same research group or author, by searching the author names in Google Scholar and downloading additional papers, the extent of cross-checking between papers for inappropriately duplicated images was naturally limited due to the large number of comparisons that could be made. No automated tools or software were used during this process.

### Screening of Papers with ImageTwin.ai

Imagetwin.ai is a commercially available AI-based software which has been developed for detecting integrity issues in scientific papers. This software can detect similar areas within images (cloned sections), between images in the same paper, and includes a database of figures sourced from open-access papers and can therefore check for images duplicated between some publications. ImageTwin.ai identifies areas of similarity, but whether the duplication is appropriate or not must be determined by the user. In this study ImageTwin.ai was used to scan the PDF files of papers that were identified as containing relevant images. Duplications highlighted by ImageTwin.ai were evaluated as appropriate or inappropriate by one reviewer (the author)

### Categories of Inappropriate Duplication

Inappropriate image duplications were assigned to three levels; approximately corresponding to previously published categories and expanded to include duplications between papers^2^:

- Category I: Simple duplication. An image is entirely reproduced without modification within the same paper, or in another paper, and labelled as representing a different experimental condition or subject.
- Category II: Duplication with repositioning. An image overlaps with another image found elsewhere in the same paper, or in another paper. The overlap is evidence that both images are smaller parts of a larger image or separate images of the same feature. The images are labelled as showing different experimental conditions or subjects.
- Category III: Duplication with alteration. Images are published with significant alteration, for example the copying or cloning of parts of an image, moving or rotating elements of a gel.

Examples of category I, II, and III duplications as identified in *Toxicology Reports* are shown in Figure 1 A, B, and C, respectively. Where multiple categories of duplication were present in one paper, the highest category was assigned to that paper. Newly Identified duplications were posted on the website PubPeer, with a short explanation and a request for the authors to respond, the corresponding author’s email was submitted to the website (which is designed to generate a system email informing the author of the comment). When a duplication had already been identified and posted on PubPeer, this was added to the dataset and classified as above.

**Figure 1:**
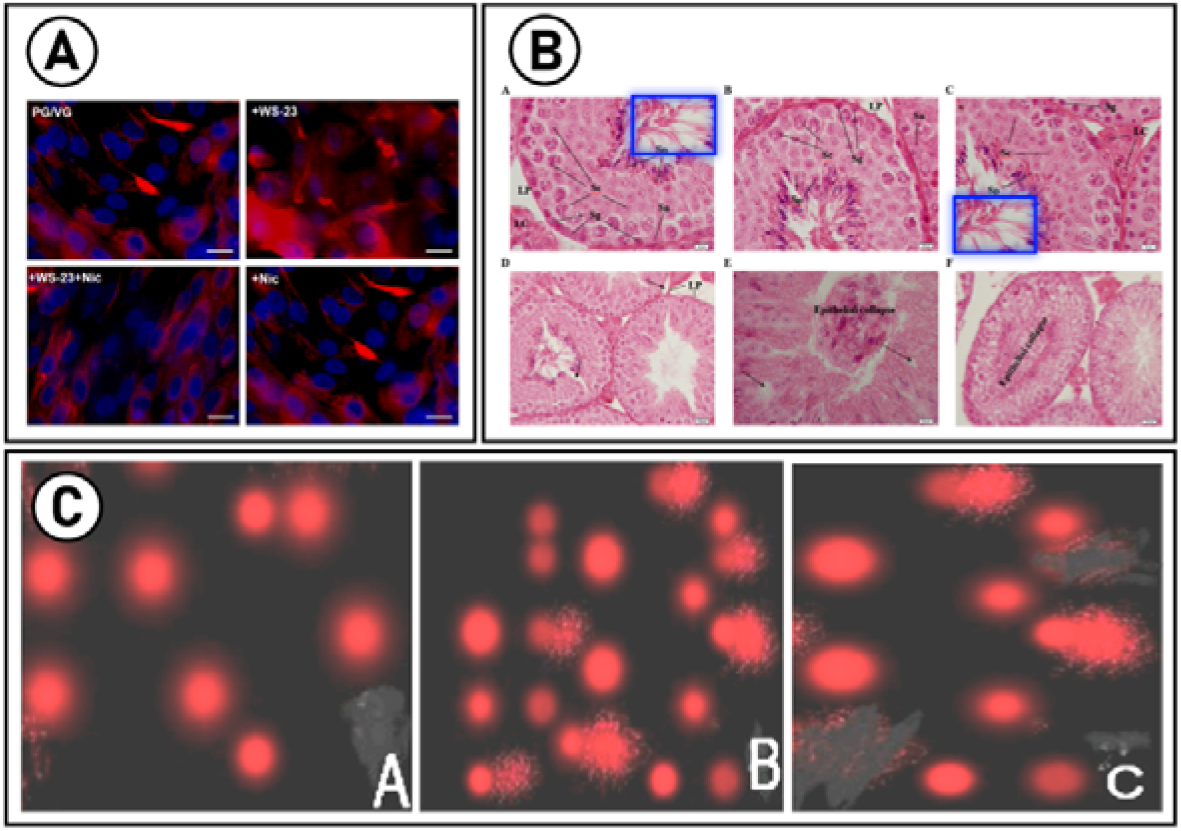
Selected examples of category I, II, and III duplications identified in Toxicology Reports. The figures have been reproduced without removing the original labels, for clarity the letters in black circles are used to identify each part of the figure. (A) Is a category I duplication, the images of cells labelled as “PG/VG” and “+Nic” in the top left and bottom right are identical. (B) Is a category II duplication, these histology images were labelled as different experimental conditions, but share a common area (blue rectangles). (C) Is a category III duplication, the central image and right-hand image share a common source (right-hand image is stretched and magnified in comparison). Parts of each image have been digitally edited with a paintbrush type tool, the brightness and contrast of (C) was increased to help visualize the edited areas.

### Comparison between Manual and ImageTwin.ai Assisted Screening

Each paper that was found to contain inappropriate duplications by the manual or ImageTwin.ai assisted screen was assigned to one of three categories:

- At least one inappropriate duplication was identified during the manual review, and none were highlighted by the ImageTwin.ai software.
- At least one inappropriate duplication was identified by ImageTwin.ai, and none were found during the manual screen.
- At least one inappropriate duplication was identified during the manual screen, and at least one during the ImageTwin.ai assisted screen. The detected duplications may be in the same images or figures, or separate.

Image duplications that had already been submitted to PubPeer by other users were excluded from the comparison between manual and ImageTwin.ai assisted screen since the method used to detect these duplications is unknown.

## Results

### Number of Inappropriate Duplications and Categories

From 2014 to the time of writing, *Toxicology Reports* published 1540 unique articles (identified by DOI), 1418 were classified as research papers or case reports during the initial screen, with 122 being reviews, retraction notices, letters, commentaries, etc. Of the research articles, around half (715) used images to illustrate the results or subject of the paper, with the other 703 papers using only charts, tables, and diagrams. Of those papers that included at least one relevant image, 115 included at least one inappropriate duplication (16%). Out of these inappropriate duplications 34 were classified as category I, 57 as category II, and 24 as category III. A Sankey diagram showing the flow of papers through this study, and the categorisation of inappropriate duplications is shown in Figure 2.

**Figure 2:**
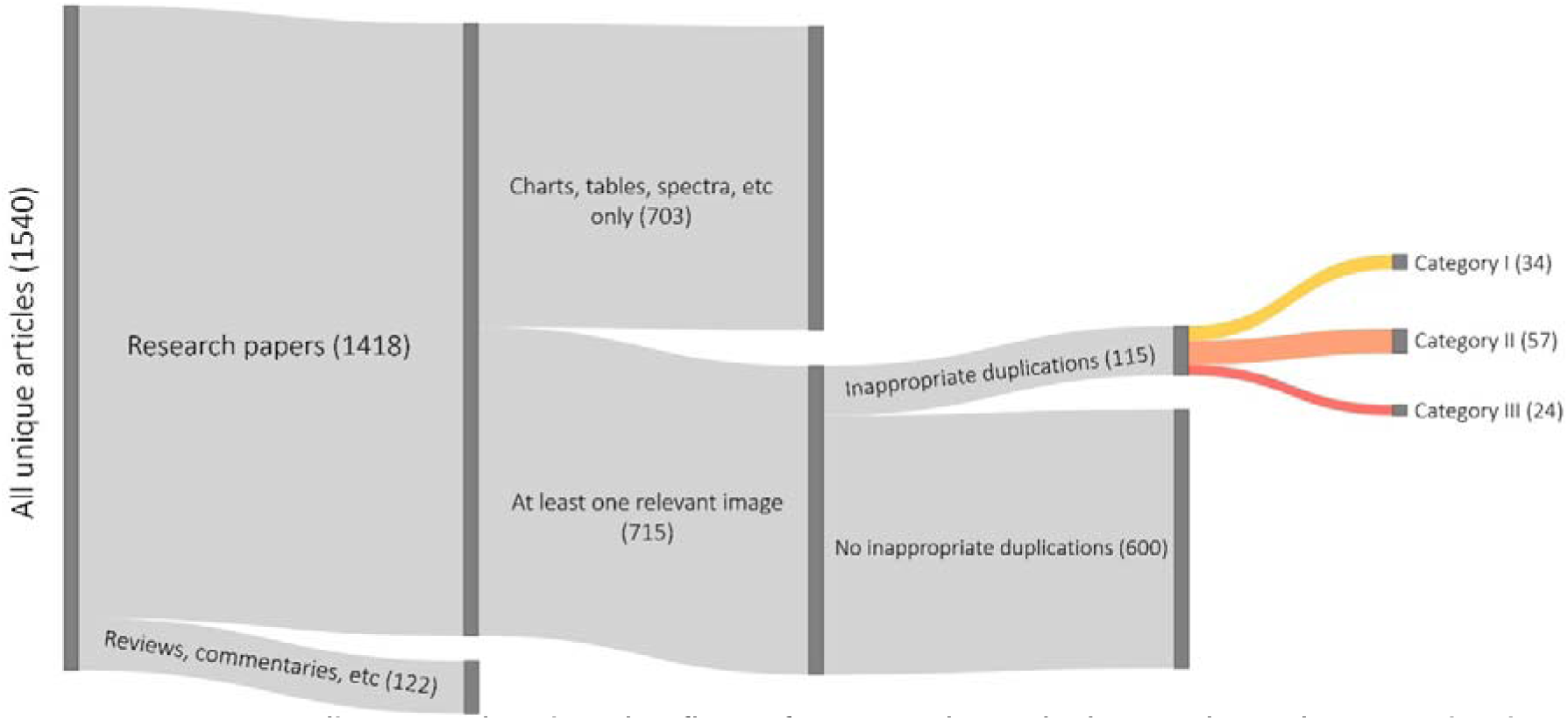
Sankey diagram showing the flow of papers through the study and categorisation of duplications. The width of each bar is proportional to the number of papers in each section.

### Comparison of Manual Screen to ImageTwin.ai Assisted Screen

A total of 63 papers containing inappropriate duplications were detected during the manual screen, and subsequently the ImageTwin.ai assisted screen identified a further 41, Prior to the start of this study 11 papers containing inappropriate duplications had been posted to PubPeer by other users, for a total of 115 papers, Figure 3 shows a Venn diagram of these results.

**Figure 3:**
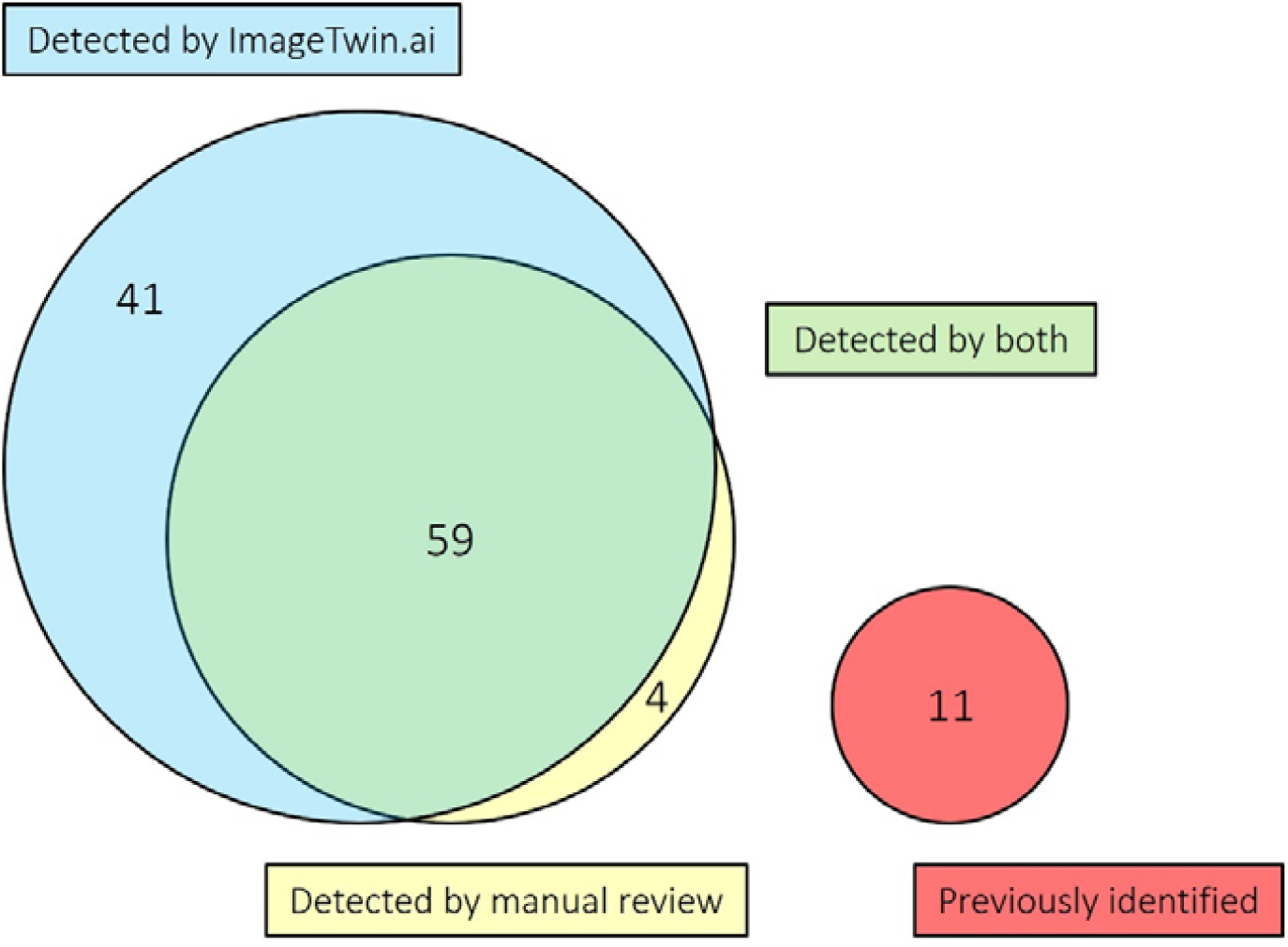
Venn diagram showing how papers containing problematic duplications were detected. Eleven papers were previously identified by unknown methods and posted to the PubPeer website.

## Discussion

### Number of Inappropriate Duplications and Categorisation

When considering only those articles containing relevant images, the rate of inappropriate duplication in *Toxicology Reports* has been quantified at 16%, if all research papers are counted (regardless of whether the paper contained any images) the rate is 8%. These estimates align with previous work^1,2,5^. About 30% of problematic papers contained only category I duplications, this type of duplication may represent a simple mistake. Around 50% of papers contained at least one category II duplication, this type of duplication is seen as more likely to be indicative of research misconduct. About 20% of papers contained category III duplications, with significant alterations of an image, including cloning of areas.

### Comparison Between Manual Review and Screening Assisted by ImageTwin.ai

The use of ImageTwin.ai increased the detection of papers containing problematic duplications. This comparison is admittedly limited, as different reviewers may have detected more or less duplications at the manual review stage. Of the four images that were identified only by manual screening and missed by ImageTwin.ai, two were category II duplications and two were category III. More extensively manipulated images may not be identified by the software, but can still be spotted by eye, the example in Figure 1C was not identified by ImageTwin.ai. Even though ImageTwin.ai was able to increase the number of duplications identified it is worth noting that the majority were spotted by one reviewer and could have presumably been identified at the peer review stage.

### Risk of False Positives and False Negatives

The true number of inappropriately duplicated images published in *Toxicology Reports* remains unknown. False negatives (duplications exist but were not identified) are likely to exist in this dataset due to the failure of the reviewer and software to identify matching areas. False positives (an image was categorised as an inappropriate duplication when it is not) may also exist within the dataset. One false positive was previously identified by an author responding on PubPeer, and this example was removed from the dataset prior to analysis. The risk of false positives and difficulties in determining the true number of duplications should not discourage research into scientific misconduct, encouragingly, three corrigenda have been published by *Toxicology Reports* which were initiated by the authors after corresponding comments were posted to PubPeer^6–8^. Alternative approaches, for example generating synthetic datasets containing manipulated images will likely prove useful to benchmark the performance of automated tools^9^.

## Conclusion

The rate of Inappropriate duplication of images in the journal *Toxicology Reports* is surprisingly high. This study adds a further quantitative result to research on scientific misconduct. Furthermore, a real-world comparison between manual review and AI assisted screening has been reported. Future research could benefit from reviewing a more diverse set of papers, expanding the types of image manipulation considered, the participation of second or third reviewers, and evaluation or comparison of different software.

## Supporting information

supplementary data set

